# Influence of High-Fat Diet and Sex on Connective Tissue in Carpal Tunnel Syndrome

**DOI:** 10.1101/2023.07.15.549152

**Authors:** Tomoyuki Kuroiwa, Hayman Lui, Koichi Nakagawa, Naoya Iida, Claire Desrochers, Rou Wan, Ken Nishimura, Elameen Adam, Dirk Larson, Peter Amadio, Anne Gingery

## Abstract

Carpal tunnel syndrome (CTS) is a common musculoskeletal disorder, characterized by thickening and fibrosis of the subsynovial connective tissue (SSCT). Risk factors for CTS includes sex, metabolic dysfunction and age. In this study we hypothesized that a high-fat diet (HFD), a common driver of metabolic dysfunction, would promote SSCT thickening in CTS and that this response would be sex dependent. To test this, we examined the effects of HFD and sex on SSCT thickening and markers of fibrosis using our established CTS rabbit model of SSCT thickening. Forty-eight (24 male, 24 female) adult rabbits were split into four groups including HFD or standard diet with and without CTS induction. SSCT was collected for histological and gene expression analysis. HFD promoted SSCT thickening and upregulated profibrotic genes, including TGF-β. Fibrotic genes were differentially expressed in males and females. Interestingly while the overall prevalence of CTS is greater in women than in men, under conditions of metabolic dysfunction men have a higher incidence. This suggests a focus on metabolic and sex specific therapeutic strategies for the treatment of patients with CTS.

## Introduction

Carpal Tunnel Syndrome (CTS) is an idiopathic non-inflammatory fibrotic age-related disorder in which progressive fibrosis of the subsynovial connective tissue within the carpal tunnel, is associated with progressive compression and dysfunction of the median nerve. Despite this prevalence, in most cases the underlying cause of CTS is unknown. Surgery is the only effective curative treatment ^1^. A better understanding of the etiology of CTS is important, as this may lead to new nonsurgical treatments, targeting the underlying mechanism of disease.

Fibrosis and subsequent thickening of the subsynovial connective tissue (SSCT) is driven by transforming growth factor beta (TGF-β)-mediated signaling ^2-4^. A significant risk factor for CTS is metabolic dysfunction^5^,which has been proposed to induce systemic inflammation that promotes fibrosis in various tissues and organs ^6^. Multiple studies have reported that metabolic dysfunction and high fat diet (HFD) promote TGF-β-mediated fibrosis ^7-14^.

Another consideration when evaluating CTS etiology is the sex difference in the prevalence of CTS, which occurs twice as frequently in women as in men^15^. Interestingly, men with metabolic dysfunction are more likely than women to have CTS ^16,17^. Based on this information, we hypothesized that there is a sex difference in the relationship between CTS SSCT thickening, markers of fibrosis and metabolic dysfunction. We also hypothesized that HFD ^18-20^ which drives metabolic dysfunction and induces SSCT thickening in the carpal tunnel in a sex dependent manner. To test this hypothesis, we conducted a comparative study to evaluate SSCT thickening and markers of fibrosis in mature male and female rabbits who were fed a HFD or standard normal diet (STD), with or without CTS fibrosis induction (Figure 1). ^21-23^

**Figure 1.**
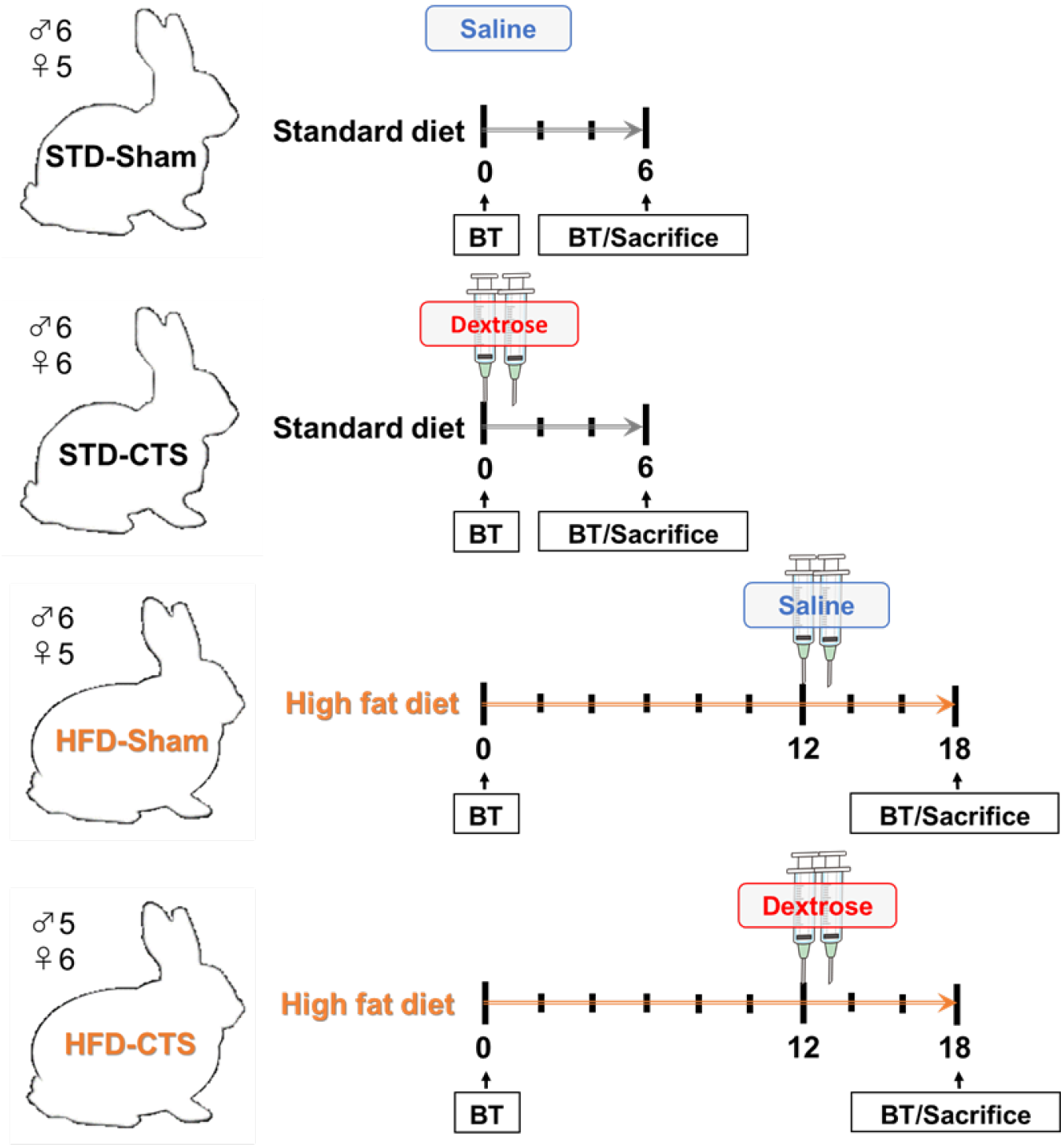
Flow diagram of the study. Rabbits were divided into four groups - high fat diet (HFD) or standard diet (STD) and CTS induction (CTS) or Sham. Rabbits were sacrificed 6 weeks after the first CTS dextrose injection. Blood tests (BT) were performed at the time of sacrifice.

## Results

### Metabolic dysfunction

Blood analyses were completed at sacrifice. The results showed no significant interactions between diet and sex (Figure 2). HFD female and male rabbits had significantly higher cholesterol, triglycerides, and insulin levels and significant lower glucose levels as compared to STD rabbits (Figure 2).

**Figure 2.**
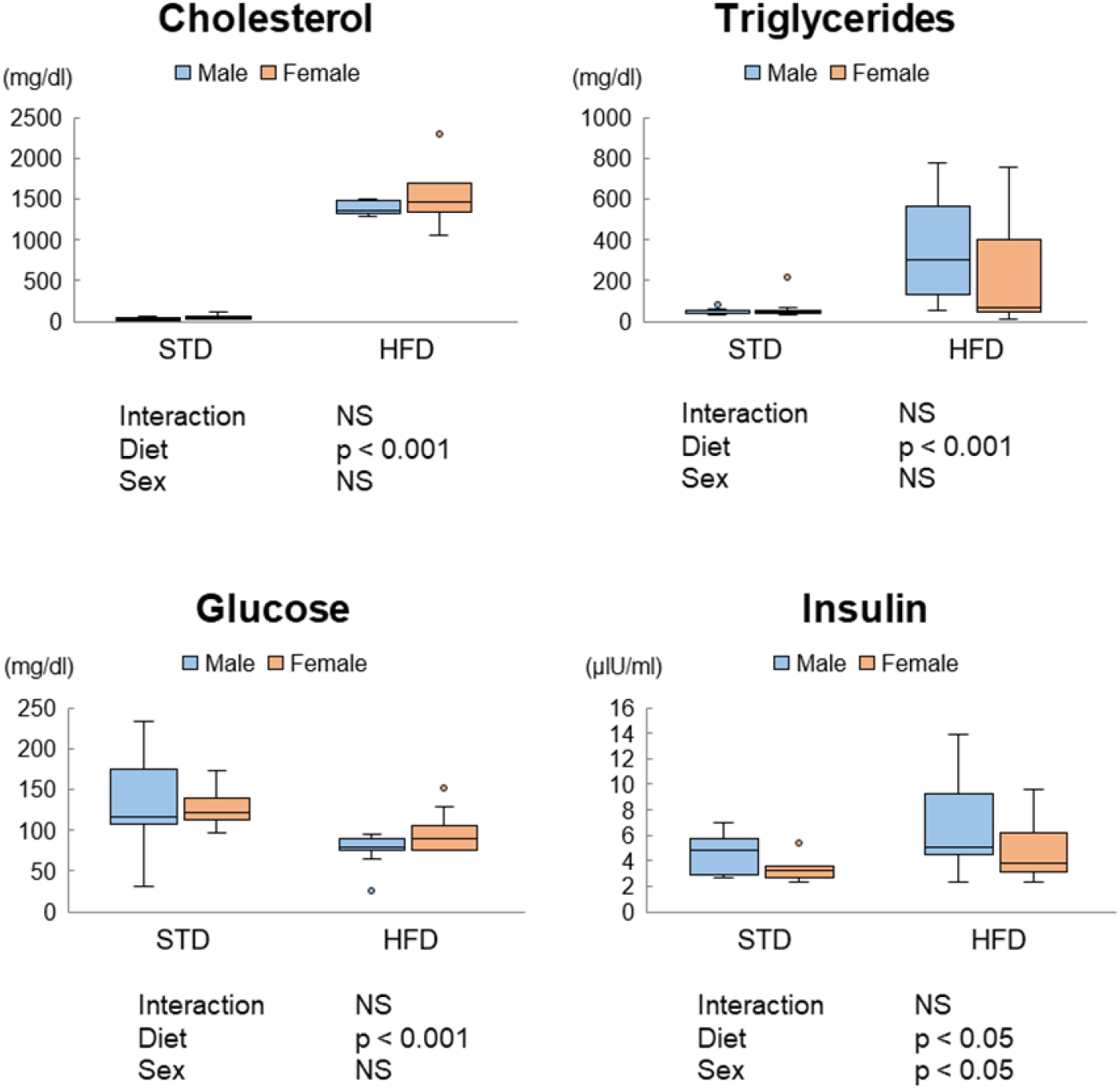
Blood analysis. Cholesterol, triglycerides, glucose, and insulin levels of rabbits at sacrifice were measured. Diet significantly regulated cholesterol, triglycerides, glucose, and insulin levels. Significant sex differences were found in insulin levels. No significant interactions were found between diet (HFD or STD) and sex. ‘P values for the main or interaction comparisons (diet and sex) are shown. NS indicates not significant. n=minimum 11 per group.

### SSCT histology

SSCT thickness was measured (Figure 3). A representative image of hematoxylin-eosin SSCT measurement is shown in Figure 3a. The overall average SSCT thickness was 0.16 mm, 0.31 mm, 0.33 mm, and 0.33 mm in the STD+Sham, STD+CTS, HFD+Sham, and HFD+CTS groups, respectively. The analysis showed that there was a significant interaction between diet and CTS induction and that both main effects of induction and diet were significant in affecting SSCT thickness (Figure 3b). In male rabbits, both diet and CTS induction resulted in significantly thicker SSCT (Figure 3c). In contrast, CTS induction alone in female rabbits was found to be the main driver of SSCT fibrosis. (Figure 3d).

**Figure 3.**
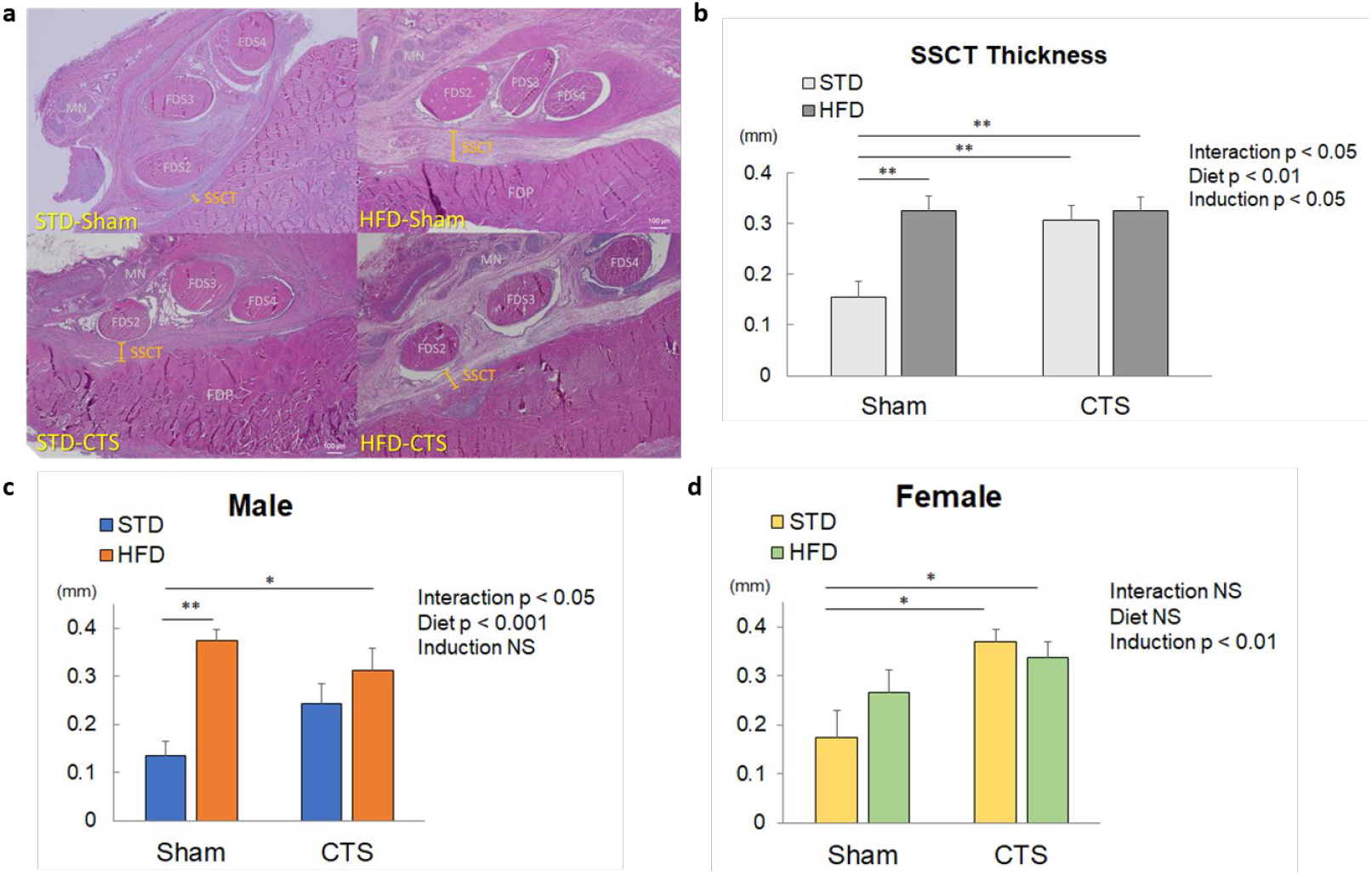
SSCT Thickness. (a) Representative histological cross-sectional images of the carpal tunnel inlet (H&E staining) in STD+Sham and HFD+Sham groups. Subsynovial connective tissues (SSCT) thickness between flexor digitorum superficialis (FDS2) and flexor digitorum profundus (FDP). (b) Average SSCT thickness in CTS and Sham induced group treated with high fat diet (HFD) or standard diet (STD). n=11. The interaction between diet and CTS induction was significant. (c) Assessment by sex showed that in male rabbits there were significant interaction between diet and CTS induction. (d). Female rabbits CTS induction was the primary driver of SSCT thickness. Means ± standard errors are depicted. P values for the main or interaction comparisons (CTS induction and diet) are shown. *, and ** indicate significant differences between the respective groups with p value < 0.05, and < 0.01, respectively. NS indicates not significant, (n = 5-6 per group).

### SSCT Gene expression

Fibrotic gene expression differences between CTS treatments were reported in scatter plots on CTS fibrosis induction or sham and diet. High fat diet (HFD) and standard diet (STD) were assessed Figure 4a-d. Multiple fibrosis-promoting genes, including Itga1, Itgb1, Tgfbr1, Ilk, Stat6, and Vegfa were upregulated in the combination of CTS induction and HFD. Significant main effects of induction and diet were found in 45 and 17 of the 84 fibrotic genes assessed, respectively (Table S-2). Fibrosis-promoting genes, including Il1a, Itgav, Tgfb3, Tgfbr1, Timp1, and Smad7 were upregulated by CTS induction alone. Furthermore, it was observed that HFD fibrosis-promoting genes including Ctgf and Cxcr4 were upregulated, and genes such as Dcn and Ifng were down-regulated in HFD rabbits. (Figure 4e and Table S-2).

**Figure 4.**
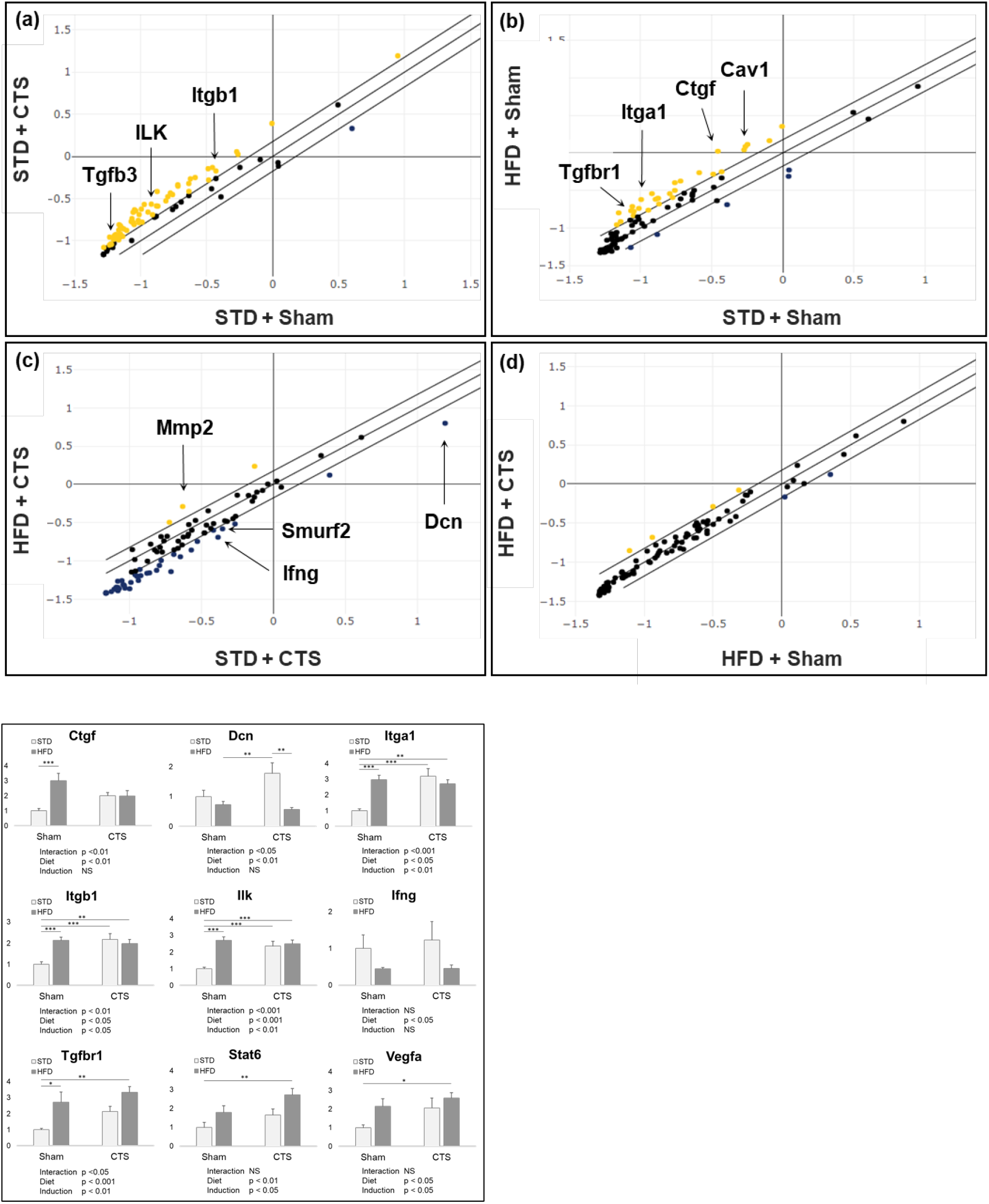
Fibrotic gene expression differences between CTS treatments were reported in scatter plots comparing CTS fibrosis induction or sham and diet for both male and female rabbits. Scatter plots (a-d) Yellow dots indicate genes that showed a significant 1.5-fold or greater change relative to STD+Sham, HFD+Sham, or STD+CTS group. Blue dots indicate genes that showed a 0.5-fold or less change. P < 0.05 was considered a significant difference, (n = 5-6 per group). (e) Fold gene expression as compared to sham controls for select genes. Expression values were normalized to STD+Sham. Interaction indicates the significance of the interaction between CTS induction and diet. Means ± standard errors are depicted. P values for the main or interaction comparisons (CTS induction and diet) are shown. *, and ** indicate significant differences between the respective groups.

In sex subgroup analysis, it was observed that the combination of male sex and a HFD resulted in 8 significantly upregulated genes, including Cul1, Cxcr4, Eng, Fli1, Prkcd, Stat6, Tgfbr1, and Timp4 (Table S-3). In contrast, female interaction between sex and diet demonstrated only one upregulated gene (Illb) (Table S-3). No down-regulated genes were observed in HFD in both sexes (Table S-3). In males, significant main effects of CTS induction were found in 28 genes and in diet 3 regulated genes. In females, significant main effects of CTS induction were found in 4 genes and in diet 13 genes. Of the 46 genes that demonstrated significant main effects, only two genes had the same main effect in both males and females (Itgav and Tgfb3)(Figure 5 and Table S-3).

**Figure 5.**
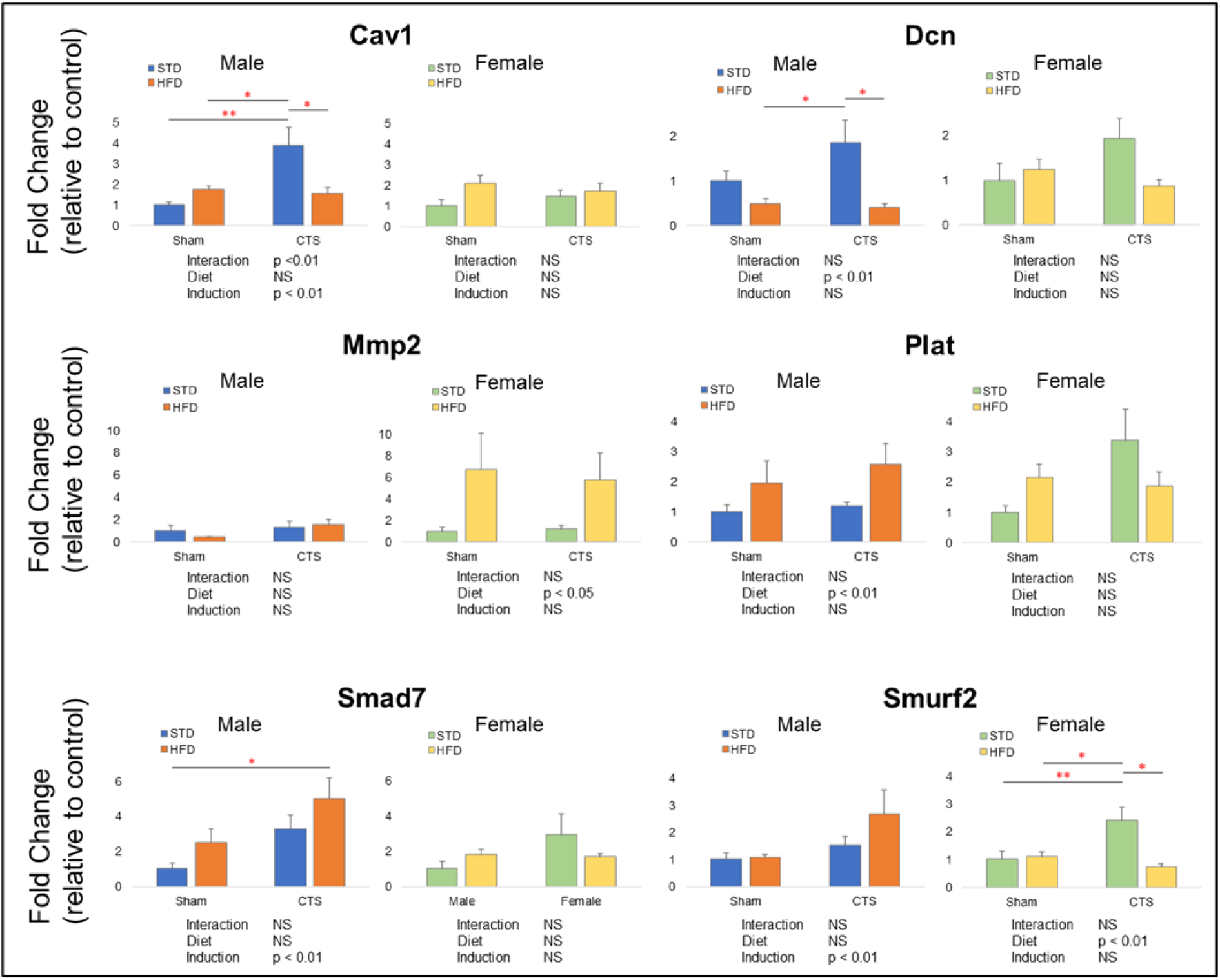
SSCT gene expression of CTS induction and diet for females and males Fold changes normalized to male STD+Sham or female STD+Sham groups. Induction and diet indicate the significance of the main effects of the CTS induction and HFD, respectively. Means ± standard errors are depicted. P values for the main or interaction comparisons (CTS induction and diet) are shown. *, and ** indicate significant differences between the respective groups.

## Discussion

CTS is an idiopathic non-inflammatory fibrotic age-related disorder in which progressive thickening of the subsynovial connective tissue within the carpal tunnel can result in the progressive compression and dysfunction of the median nerve. Incidence of CTS is associated with metabolic dysfunction, age and sex^15-17^.

In this study a rabbit model of CTS thickening and HFD were used as previously described ^22,24-27^. All rabbits exhibited induced hypercholesterolemia with HFD. CTS induction and HFD upregulated many fibrosis-inducing genes including Tgfbr1, a receptor protein of the TGF-β family, as well as fibrosis-associated integrin-related genes (Itga1, Itgb1, Ilk) ^2,28-30^, Stat 6 ^31^, and Vegfa^32,33^. Additionally CTS fibrosis induction and HFD significantly increased SSCT thickness.

Since CTS and metabolic dysfunction is more prevalent in middle-aged patients, we selected mature animals to more accurately represent the clinical condition studied. HFD treated rabbits used in this study developed hypercholesterolemia and hypertriglyceridemia similar to previous reports^22^. In contrast we did not find a significant increase in blood glucose levels. This may have been due to age differences as the present study used middle-aged rabbits compared to young rabbits in previous studies. This finding that HFD blood glucose levels are not significantly increased in mature rabbits is consistent with trends observed in other HFD-fed animal studies using both young and aged animals which also report a significant elevation in blood glucose levels only in young animals only ^34,35^.

It is well established that metabolic dysfunction promotes fibrosis in many tissues/organs ^6,36^. Clinically, metabolic dysfunction has been reported as one of the risk factors for CTS^5,37-39^, and the results reported here provide support for HFD driving SSCT thickening. Interestingly, subgroup analysis of sex showed that the HFD induced SSCT thickening in male rabbits only and that the regulated gene profiles are different between treatments. Similarly previous studies have reported that mature male rats and mice were also more vulnerable to the effects of the HFD than females ^40,41^. Our findings are consistent with clinical studies have reported that metabolic dysfunction is a risk factor for CTS ^5,37-39,42^, and that this incidence is higher in males^16,17^.

We and others have shown that Ctgf is a known mediator of TGF-β induced fibrosis ^4,43-45^. In this study, HFD resulted in significant increases in Ctgf. This response has been observed in previous HFD animal studies^46-48^ and in analysis of SSCT tissue of patients with CTS^2,3^. Additionally, Tgfbr1 and integrin-related genes were observed to be significantly upregulated in both CTS induction and HFD fed animals. Tgfbr1 encodes a receptor that binds to TGF-β protein, promotes TGF-β signaling, and has also been found to be upregulated with metabolic syndrome ^4,12,44,45,49,50^. A subset of the integrin family has been reported to play an important role in the activation of latent TGFB1, thereby promoting the fibrosis process in models of various organ fibrosis ^29,30,51,52^. The significant upregulation of these genes is consistent with the results of the SSCT thickness assessment.

In contrast, the expression of anti-fibrotic genes such as Ifng^53,54^ and Dcn^55-57^ were noted to be significantly downregulated in HFD rabbits. Interferon-γ, encoded by Ifng, is an immune-regulatory cytokine that also serves as a strong anti-fibrotic agent^54^. IFN-γ also inhibits fibroblast proliferation and collagen deposition and downregulates Tgfb expression^53^. The combination of both upregulation of fibrotic genes and downregulation of anti-fibrotic genes in HFD rabbits suggests that metabolic dysfunction induces gene expression changes that may drive SSCT fibrosis.

Gene expression levels also differed between males and females. In male rabbits, CTS induction significantly upregulated Cav1 and Smad7. Cav1 is overexpressed in adipose tissue of obese patients and is positively correlated with body fat and body mass index^58^. Cav1 has also been reported to suppress fibrosis in various organs via suppression of TGF-β/Smad signaling ^59,60^. Smad7, a Smad inhibitory protein, is a key regulator that binds to regulatory SMADs and acts in to inhibit TGF-β signaling. Smad7 overexpression has been shown to antagonize TGF-β-mediated fibrosis and inflammation ^61,62^. Therefore, the observed elevation in Cav1 and Smad7 expression in male rabbits may be a compensatory response to ongoing fibrosis in the SSCT.

Furthermore, Dcn, another negative regulator of TGF-β signaling, was significantly down regulated in HFD males only. Decorin, a small leucine-rich proteoglycan encoded by Dcn, regulates collagen fibrillogenesis during development. It exerts a protective effect against fibrosis in various tissues by directly blocking the bioactivity of Tgfb1^55-57^. The loss of this important TGF-β regulator in conditions of metabolic dysfunction may contribute to increased SSCT fibrosis in males.

Another gene that was regulated by diet in males was plasminogen activator encoded by Plat. Plat is a component of the plasminogen system that is important for proteolytic remodeling of extracellular matrix. Plasminogen activators can act to attenuate the action of plasminogen activator inhibitors (PAI-1) that can drive fibrosis and extracellular matrix deposition ^63^. Expression of Plat could be a compensatory mechanism to attenuate fibrosis. Plat also has been shown to limit activation of Tgfb1^64,65^.

In contrast, HFD significantly upregulated Mmp2 and downregulated Smad ubiquitin regulatory factor 2 (Smurf2) in females only. MMP2 has been shown to have anti-fibrotic effects^66,67^ and Smurf2 negatively regulates TGF-β signaling by ubiquitinating TGF-β receptors (TβR) and Smad proteins. Together, these significant disparities in gene expression may provide a mechanistic explanation for the disparate response to CTS induction and diet in males and females.

In this study, female rabbits were housed two to a cage, whereas male rabbits were housed 1 to a cage, as required by our IACUC. This difference in housing may have caused differences in exercise and other behaviors. HFD rabbits were maintained on HFD for 12 weeks, in contrast the CTS or sham-induced STD rabbits which were treated after the standard 1 week of acclimation for all animals. The effect of the difference in length of housing in our facility is likely to be negligible, as the duration of housing in our facility for the treatment phase was identical in all four groups. Our future work includes assessment of sex differences in the gene and protein regulation of Cav1, Dcn, Mmp2, Plat, Smad7, Smurf2 in human SSCT biopsies.

In summary, the effect of HFD and its interaction with sex on SSCT thickening in a CTS animal model was studied. HFD resulted in increased fibrotic gene expression and SSCT thickness compared to STD rabbits. This is consistent with previous analysis of the SSCT in CTS patients^23,68^. In addition, the combination of HFD and CTS induction in males resulted in significantly greater SSCT thickening when compared to females. This is supportive of previous reports indicating that men with metabolic dysfunction are more likely to develop CTS ^16,17^. For the first time, this study has shown a potential mechanism of the differential sex-based development of CTS in patients with metabolic dysfunction.

## Material and Methods

### Animals

This study was approved by Mayo Clinic Institutional Animal Care and Use Committee (IACUC A5660), and all procedures were performed in accordance with the National Institutes of Health Guide for the Care and Use of Laboratory Animals. The CTS SSCT fibrosis rabbit model was created by our group and is the smallest animal with human-like SSCT ^23,27,68,69^. This model recapitulates the SSCT thickening and fibrosis found in CTS patients. 48 retired breeder New Zealand White rabbits (24 females, 24 males) were used to represent middle-aged animals, as CTS is an age-related disease with its peak incidence in humans aged 45-55. For this reason, we used mature rabbits with an average age of 2.6 (0.9 - 4.3) years.

Rabbits were randomly divided into four groups of 12 (6 females and 6 males in each group). The groups were fed a standard diet (STD) or a high-fat diet (HFD) and SSCT fibrosis was induced by dextrose injections which drives SSCT thickening (CTS) or saline injections (Sham) as previously described ^27^. Groups were designated as follows: STD+Sham, STD+CTS, HFD+Sham, and HFD+CTS (Figure 1). Rabbits in the STD groups were acclimated for a week on STD and then received the CTS or Sham injections as described below. STD rabbits continued the STD for an additional 6 weeks prior to sacrifice. HFD group rabbits were acclimated for a week and fed HFD for 12 weeks prior to injections to develop metabolic dysfunction, based on previous studies of HFD in rabbits ^24-26^. The HFD rabbits then received CTS or Sham injections and continued the HFD for an additional 6 weeks prior to sacrifice. (Figure 1). In all groups, immediately after sacrifice, one forepaw was harvested for histology and the other for SSCT gene expression analysis. Metabolic dysfunction was assessed by measuring hyperglycemia, hypercholesterolemia, and hypertriglyceridemia dysfunction as described below ^70^. One STD-fed female and one HFD-fed female rabbit died during the study. The former died from causes unrelated to the study design, whereas the latter died from liver failure, associated with the HFD.

### Diet

The diet for the STD group was a standard rabbit diet (Laboratory Rabbit Diet HF 5326, CA). This diet included 12% water, 14.5% protein, 23% fiber, and 3.8% fat. The HFD (Teklad Custom Diet TD.200785, ENVIGO, IN) was the STD supplemented with 0.5% cholesterol and 4% peanut oil, as previously reported ^21,22,24,25^. Water and food were not restricted during this study.

### CTS Induction

Using our established rabbit model of CTS, a small incision was made with direct vision while under anesthesia. For CTS and sham, the carpal tunnels of both forepaws were injected with either 0.1 ml of 10% dextrose solution (CTS) or 0.1 ml of 0.9% saline (Irrigation, Baxter, IL) (Sham), using a 30-gauge needle. As per the protocol two injections were given 1 week apart. Each injection was made inside the synovium around the middle digit flexor digitorum superficialis tendon, as described previously ^23,68^.

### Metabolic Dysfunction Assessment

Blood samples were taken at sacrifice. Rabbits were fasted overnight and blood was taken from the marginal ear vein. Blood samples were centrifuged at 3,000 rpm for 20 minutes at 4 °C, and the supernatant was collected as serum. Samples were sent to IDEXX BioAnalytics (Columbia, MO) for analysis of total cholesterol, triglyceride, glucose, and insulin levels in the serum (IDEXX BioAnalytics Case # 82497-2021).

### Histology

The tissue was harvested so that an axial section at the wrist level was visible after bone removal. The SSCT was fixed in a 4% paraformaldehyde solution for approximately one week, embedded in paraffin, and sectioned into 4-μm-thick slices. Sections were stained with hematoxylin-eosin and observed under a light microscope (All-in-One Fluorescence Microscope BZ-X800, Keyence, IL, RRID:SCR_019119).

### Image analysis

Tissue images were evaluated using ImageJ software (ImageJ, National Institutes of Health; http://imagej.nih.gov/ij/, RRID: SCR_003070) by double blind observers. The distance between the flexor digitorum superficialis (FDS) and flexor digitorum profundus (FDP) was measured and determined to be the SSCT thickness, as described in previous studies ^23,71^ (Figure 3). The average values of the thickness measured by two observers were used for analysis.

### Quantitative real-time polymerase chain reaction (PCR)

After sacrifice, total RNA was extracted from the SSCT of one forepaw using Trizol reagent (Invitrogen Life Technologies, NY). The cDNA was obtained from equal amounts of total RNA by reverse transcription using an iScriptTM cDNA Synthesis Kit (Bio-Rad, CA). Gene expression of 96 fibrosis-associated genes were evaluated using a PCR array kit (RT^2^ Profiler™ PCR Array Rabbit Fibrosis PANZ-120Z, Qiagen) (Genes are listed in Table S-1) by quantitative real-time PCR using a Thermal Cycler (Bio-Rad Laboratories, CA, RRID:SCR_019152). The average values of the results in triplicate were used for analysis. Eighty-four fibrosis-related genes were evaluated. Data was normalized to the geo-mean of housekeeping genes Acta2, Actb, Gapdh, Ldha, and LOC100346936.

### Statistics

#### Sample size and power

Based on previous studies using this model, a sample size of 6 rabbits per group is required to achieve 80% power to detect a difference in means of 0.15 mm between groups in the FDS-FDP distance, assuming an overall within group standard deviation of 0.08 mm^23^. Additionally, to perform separate analyses within each sex, each group was fully powered with 6 males and 6 females, for a total of 12 rabbits per group (48 rabbits total).

#### Analysis

Separate analyses were performed for males and females. Within each sex subset, the effects of CTS induction (no vs yes) and diet (HFD vs STD) on SSCT thickness and gene expression level was evaluated using two-way analysis of variance (ANOVA). For blood test results, mean values were calculated for each diet group and sex and compared using two-way ANOVA. Holm-Bonferroni post-doc tests were completed. P-values less than 0.05 were considered statistically significant.

## Acknowledgments

This study was supported by NIH-NIAMS R01AR076347 and Mayo Clinic.

## Data Availability

The data that supports the findings of this study are available in the methods and/or supplementary material of this article.

## Author Contribution

T. Kuroiwa, P. Amadio, and A. Gingery formulated the concept and designed the research; T. Kuroiwa, A. Gingery, H. Lui, K. Nakagawa, R. Wan, and E. Adam performed the experiments; T. Kuroiwa, N, Iida, K. Nishimura, and C. Desrochers acquired the data; T. Kuroiwa, D. Larson, PC. Amadio and A, Gingery analyzed and interpreted the data. All authors were involved in drafting and revising the manuscript. All authors have read and approved the final submitted manuscript.

